# Molecular architecture of the autoinhibited kinesin-1 lambda particle

**DOI:** 10.1101/2022.04.28.489841

**Authors:** Johannes F. Weijman, Sathish K.N. Yadav, Katherine J. Surridge, Jessica A. Cross, Ufuk Borucu, Judith Mantell, Derek N. Woolfson, Christiane Schaffitzel, Mark P. Dodding

**Affiliations:** School of Biochemistry, University of Bristol, Biomedical Sciences Building, University Walk, BS8 1TD, UK; School of Chemistry, University of Bristol, Cantock’s Close, Bristol, BS8 1TS, UK; GW4 Facility for High-Resolution Electron Cryo-Microscopy, University of Bristol; Bristol BioDesign Institute, University of Bristol, Life Sciences Building, Tyndall Avenue, Bristol, BS8 1TQ, UK

## Abstract

Despite continuing progress in kinesin enzyme mechanochemistry and emerging understanding of the cargo recognition machinery, it is not known how these functions are coupled and controlled by the alpha-helical coiled coils encoded by a large component of kinesin protein sequences. Here, we combine computational structure prediction with single-particle negative stain electron microscopy to reveal the coiled-coil architecture of heterotetrameric kinesin-1, in its compact state. An unusual flexion in the scaffold enables folding of the complex, bringing the kinesin heavy chain-light chain interface into close apposition with a tetrameric assembly formed from the region of the molecule previously assumed to be the folding hinge. This framework for autoinhibition is required to uncover how engagement of cargo and other regulatory factors drive kinesin-1 activation.

**Summary statement:** Integration of computational structure prediction with electron microscopy reveals the coiled-coil architecture of the autoinhibited compact conformer of the microtubule motor, kinesin-1.

---

The autoinhibition of cytoskeletal motors is a fundamental regulatory mechanism that governs a vast array of cellular processes, ranging from intracellular transport to cell division and migration. These states are controlled by intramolecular or intra-complex interactions that serve to inhibit motor activity until it is required and prevent the wastage of ATP. There has been good progress in understanding the nature of the autoinhibited states of dynein and myosin family members in their higher-order functional complexes (*1–6*), but insight in the kinesin family has remained limited to isolated domains and certain critical interfaces (*7–9*).

The ubiquitous and prototypic kinesin family member, kinesin-1, exists in a predominantly heterotetrameric form, comprising two motor bearing heavy chains (KHCs) and two cargo-binding /regulatory light chains (KLCs) (*10*). As it carries out its diverse functions in the transport of vesicles, organelles, mRNAs, and multi-protein complexes, kinesin-1 transitions from a compact autoinhibited state to an extended active state. This is controlled by the binding of both cargo adaptors and microtubule associated proteins (*8, 11–20*). The compact state suppresses microtubule-dependent kinesin-1 ATPase activity because it allows a regulatory short-linear-motif (SLiM) residing near the C-terminus of the KHCs (known as the isoleucine-alanine-lysine (IAK) motif) (*7, 21–27*) to bind the *N*-terminal motor domains **(Fig. 1A)**. The KLCs also engage in a second regulatory SLiM mediated self-interaction that occludes an important cargo adaptor binding site on their tetratricopeptide repeat (TPR) domains (*8, 28, 29*) **(Fig. 1B)**.

**Fig. 1.**
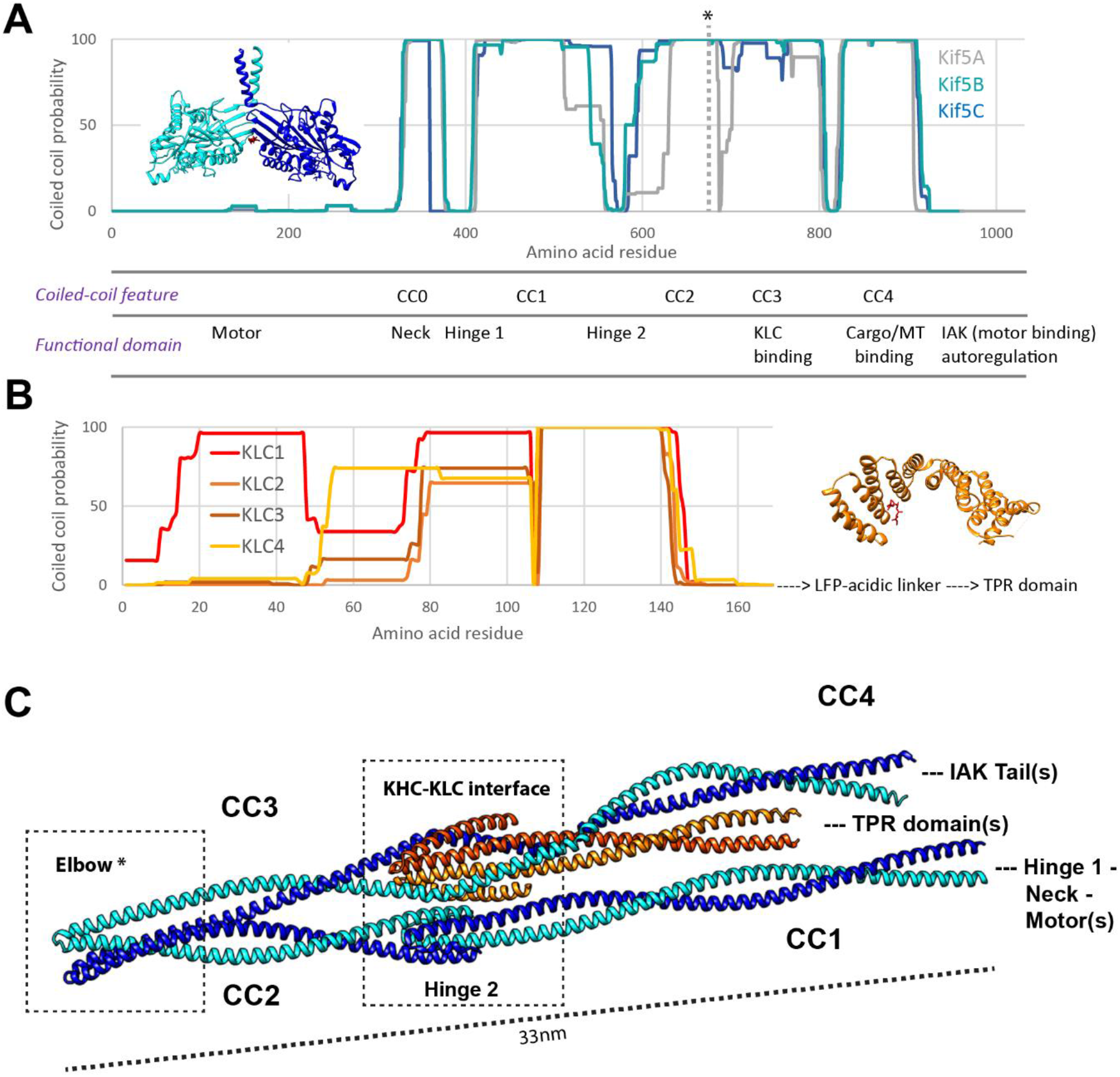
Computational prediction of a folded coiled-coil conformation for the kinesin-1 heterotetramer. (A) Coiled-coil probability plots generated using Marcoil for the mammalian KHC paralogues Kif5A-C (NP_004975.2; NP_004512.1; NP_004513.1). Inset shows crystal structure of a motor domain dimer bound a regulatory IAK peptide (Drosophila orthologue, PDB:2Y65). * on Marcoil plot indicates the position of the elbow feature. (B) Coiled-coil probability plots for the amino terminal region the mammalian KLC paralogues KLC1-4 (NP_032476.2; NP_032477.2; NP_001272967.1; NP_001275963.1). Inset shows crystal structure of TPR domain in the presence of its LFP-acidic regulatory peptide (Mouse, 5FJY). (C) AlphaFold2-Multimer prediction of a 2:2 kinesin-1 heterotetramer comprised of KHC (Kif5C, NP_001101200.1) residues S410-H917 and KLC (KLC1, NP_032476.2) residues T20-S162. Positions of the globular domains shown in A and B are highlighted.

Beyond these isolated interfaces, direct insights into architecture of the autoinhibited state have been very limited since early pioneering electron microscopy studies outlined the intrinsic flexibility of the complex and its sensitivity to ionic strength and pH (30–34). Indeed, these conformational dynamics and resulting heterogeneity of isolated complexes are likely to be the main reason why progress has been limited. As such, the organisation of the series of KHC coiled-coil domains that extend throughout the complex remains a mystery. Consequently, it is unclear how the binding of the KLC to these coiled coils helps to maintain the kinesin-1 complex in its inactive state (*12, 13, 35*).

To address these questions, we began by examining predictions of coiled-coil domains in the protein sequences of the three paralogous mammalian KHC (Kif5A, Kif5B and Kif5C) and four KLC (KLC1-4) protein sequences using Marcoil coiled coil prediction software (*36, 37*). This highlighted a series of heptad repeats indicative of coiled-coil regions in KHC, CC0 to CC4 **(Fig. 1A)**(*38*): CC0 corresponds to the neck coil; CC1-CC3 form the stalk, the latter component of which includes the KLC binding site; and CC4 is known to bind several cargoes/adaptors (*26, 39–42*). CC1 and CC2 are separated by a drop in coiled-coil prediction probability (*known as Hinge 2*) (*23, 25, 43, 44*), which has generally been thought to be the point where KHC folds in the autoinhibited state. The *N*-terminal sequence of KLC (that binds to KHC) also has heptad repeats and is predicted to form a segmented coiled coil. These are followed by an unstructured linker that leads on to the cargo binding TPR domains **(Fig. 1B)**.

To explore their structural organisation of these coiled coils, we used AlphaFold2-multimer (*45, 46*) to predict a structure for a 2:2 heterotetramer consisting of CC1 to CC4 of Kif5C and the *N*-terminal domain of KLC1 isoform A **(Fig. 1C, Fig. S1)**. These are the shortest KHC and KLC paralogues. Consistent with the Marcoil prediction, the resulting model had CC1 to CC4 in KHC. Analysis of the model using SOCKET2 (*47*) identified the signature knobs-into-holes interactions of coiled-coil structures in these domains and provided a structure-based assignment of the heptad register **(Fig. S2)**. With the exceptions noted below, the coiled-coil domains are all canonical, heptad-based, parallel dimers; though CC4 had a noticeable proline kink (**Fig. 1C, Fig. S3)**. In addition, the location of the AlphaFold2-predicted light-chain binding site in CC3 is in good agreement with biochemical data, in the absence of *a priori* structural information (*48*).

However, we note three interesting and unexpected features in this model. Firstly, the KHC-KLC interface predicts as a 6-helix bundle rather than a tetrameric coiled coil previously assumed (*49*). The bundle is a dimer of antiparallel trimeric coiled coils formed from two *N*-terminal helices of the KLC and the *C*-terminus of CC3 of KHC. Secondly, Hinge 2 falls in a parallel tetrameric coiled coil formed by the termini of CC1 and CC2, facilitated by a short helix in the loop that links them. Thirdly, these two local assemblies are brought into apposition by a flexion in the KHC dimer between CC2 and CC3. We named this new feature the *elbow*, and it is marked in Fig. 1A with *. Interestingly, comparison with the AlphaFold2 prediction of the KHC dimer without KLC **(Fig. S3)**, suggested that KLC may help to orient CC4, which is separated from CC3 by a tri-glycine (GGG) motif that kinks both assemblies but to different degrees **(Fig. S3)**. The length of the overall KHC:KLC assembly, at 33 nm long (*N*-terminus to elbow), is in accordance with the first studies from Hirokawa *et al*. that measured the distance of the flexion from the motor domains (*31*). Together these data suggest the intriguing possibility that AlphaFold2 predicts the coiled-coil assembly of the autoinhibited ground state of the complex.

To test the AlphaFold2 prediction, we isolated the full-length compact conformer of heterotetrameric kinesin-1 (Kif5C/KLC1 isoform A) from complexes expressed in *E. coli* **(Fig. 2A,B)**. Compositionally indistinguishable KHC-KLC complexes eluted in size-exclusion chromatography (SEC) in two peaks after the void, consistent with an equilibrium between two states (Peak 1 and 2, **Fig. 2B**)(*16*). To capture complexes in their post-elution state, the crosslinker bis(sulfosuccinimidyl)suberate (BS3) was added to the samples. For peak 1, this gave a major slowly migrating species on SDS-PAGE (**Fig. 2A**, band marked *) and a minor faster-migrating species (band marked **). In contrast, for peak 2, the faster-migrating species was the more abundant, with only small amounts of the slower-migrating form. Together these data suggested that proteins of peak 2 may be enriched for the compact conformer.

**Fig. 2.**
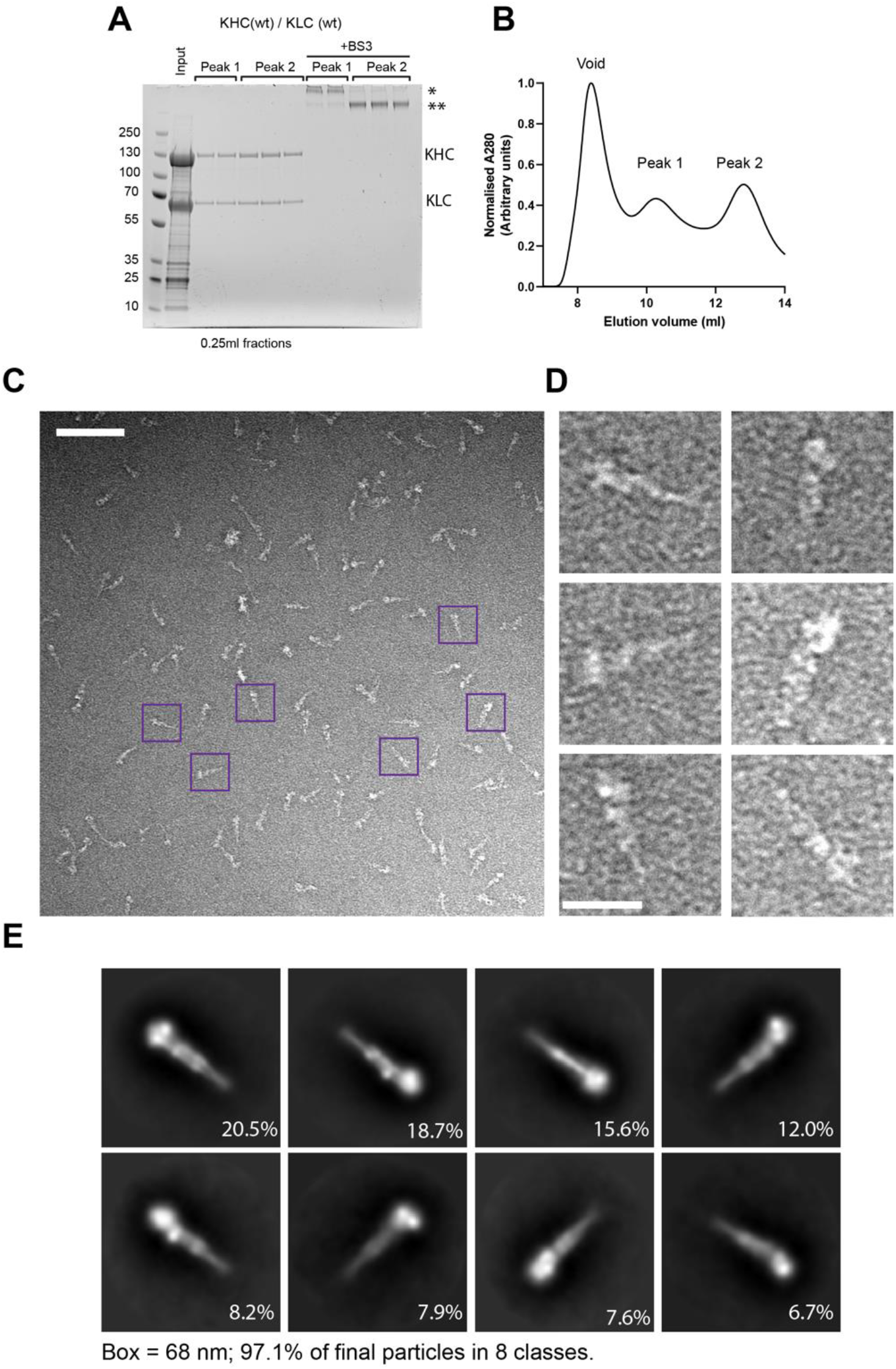
Negative stain electron microscopy reveals the architecture of the compact the kinesin-1 conformer. (A) Coomassie-stained SDS-Page gels and (B) absorbance measurements, showing SEC analysis of kinesin-1 complexes purified by nickel affinity chromatography. Complexes eluted in two distinct peaks after the void (Peak 1 and Peak 2) and 0.25ml fractions from across these peaks are shown. Lanes marked +BS3 lanes show the mobility of duplicate samples after cross-linking. Major and minor species referred to in the main text are marked with * and **. Data are representative of at least 3 independent experiments. (C) Representative T12 electron micrograph showing negative-stained, cross-linked sample from Peak 2. Scale bar is 100 nm. Purple boxed particles are expanded in (D) where the scale bar is 25 nm. (E) Reference free 2D class averages of particles from analysis of cryo-negative stain dataset.

To characterise these fractions further, we turned to negative-stain electron microscopy. We found that the major species from peak 2 was a **≈** 40 nm long V-shaped particle, with a wide end and a tapered end (82% of 279 particles counted were ≤ 45nm in length) **(Fig. 2C, D)**. The remaining particles were longer and more heterogeneous. In contrast, preparations from peak 1 revealed predominant long or bent thin particles of up to 80 nm in length—which is similar to the described open conformers (*16, 30–32*)—and relatively few of the ≈ 40 nm species (29% of 266 particles counted were ≤ 45nm length for peak 1 complexes) **(Fig. S4)**.

To study the compact species, a dataset of 3,378 micrographs was collected from peak 2 grids and processed using single particle analysis in Relion 3 (*50*). This yielded 2D class averages consistent with a single particle **(Fig. 2E, Fig. S5)**. Striking features included a pair of globular densities that, which, from their shape, size, and orientation, we attribute to the motor domains. Extending from the motor domains, there is a relatively thick rod along with additional globular densities, one close to the motor domains and one around halfway along the structure. The rod then tapers in a tight V-shape to a point. To compare the AlphaFold2 model and 2D experimental data, low-resolution back-projection images (low-pass filtered to 30 Å resolution) were generated from the AlphaFold2 model (that does not include motor and TPR domains) **(Fig. 3A, Fig. S6)**. The predicted elbow feature could be identified unambiguously. The density towards the middle of the rod is concomitant with the predicted 6-helix bundle at the KHC-KLC interface plus Hinge 2.

**Fig. 3.**
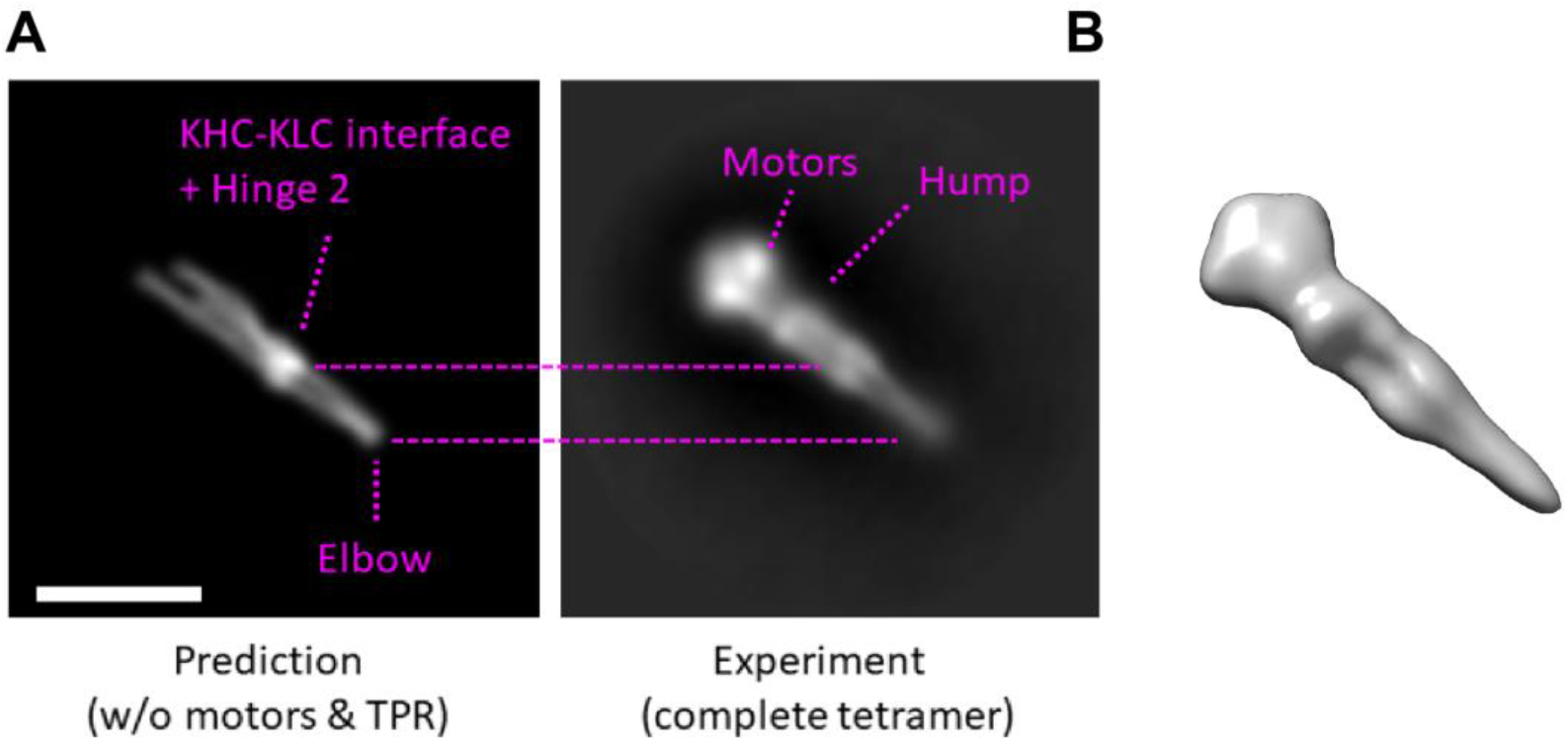
Comparison of computational and experimental data. (A) Comparison of back projection of the AlphaFold2 model (left, low pass filtered to 30 Å) with 2D class average (right) from experimental data. Boxes are 68.4 nm (B) 3D reconstruction from cryo-negative stain electron microscopy of the full length Kif5C/KLC1 tetramer from peak 2.

3D reconstruction yielded a low-resolution envelope that supported our observations from the 2D classes and was consistent with the model **(Fig. 3B)**. The position of the KLC TPR domains is less clear; we favour an interpretation where they are accommodated within the head of the complex, that is larger than would be expected for motor domains alone, and is consistent with previous fluorescence resonance energy transfer data (*12*). In which case, the hump would most likely include the AlphaFold2-predicted homodimeric coiled coil of KLC and/or CC4 of KHC that is known to be conformationally plastic, and within which, as we note above, is predicted a proline-induced kink/bulge (*39, 49*). Overall, the experimental data indicate that the computational model reflects the gross coiled-coil architecture of the compact conformer of kinesin-1.

As a further test of the model, we sought to remove the elbow and join the flanking CC2 and CC3 regions as seamlessly as possible. The aim was to open the complex and so relieve autoinhibition. Whilst the Marcoil and SOCKET2-assigned heptad repeats differ for this region (**Fig. S7**), they concur in that the regular 3-4 alternating pattern of hydrophobic residues is broken. Closer inspection identified an 18-residue stretch rich in polar charged residues, with a spacing between hydrophobics of 3-2-8, and spanning the turn predicted by AlphaFold2. Therefore, we removed this sequence to generate a Δelbow construct with a contiguous 3-4 hydrophobic spacing to join CC2 and CC3 **(Fig. S7)**. Consistent with this, kinesin-1 complexes with Δelbow eluted at the same position as wildtype peak 1 **(Fig. 4A,B)** and negative-stain electron microscopy revealed almost exclusively extended particles **≈** 80 nm long (only 2% of 229 particles counted were ≤ 45nm length) **(Fig. 4C)**. Thus, the elbow is required to form the compact conformer. Finally, consistent with motor activation, HA-tagged Kif5C with Δelbow accumulated in the cell periphery, like complexes lacking the regulatory tail (Δtail) that cannot form the compact conformer (*27*), rather than displaying the diffuse cytosolic localisation that is characteristic of wildtype complexes **(Fig. 4D)**.

**Fig. 4.**
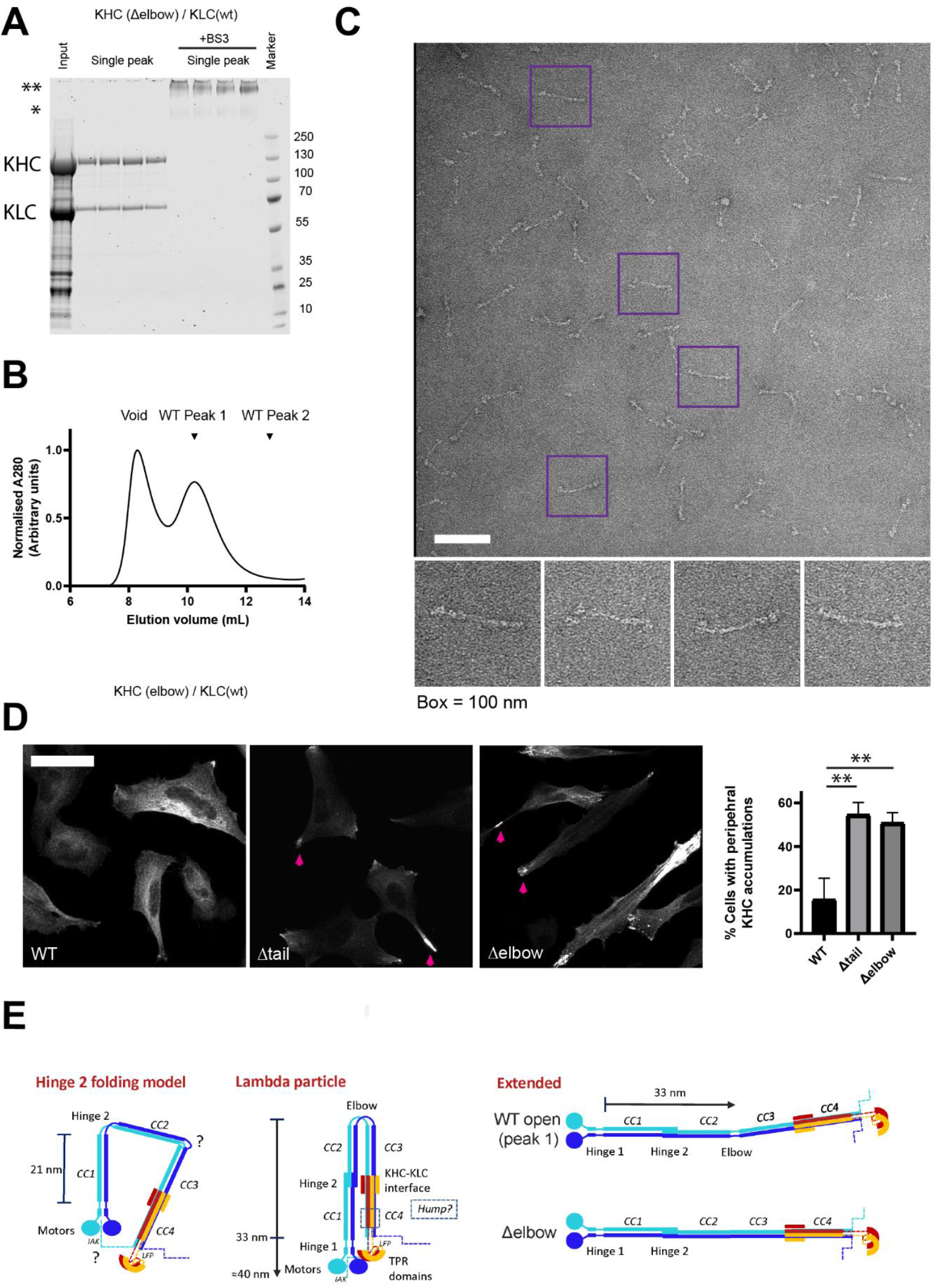
Disruption of the elbow prevents formation of the compact kinesin-1 lambda particle. (A) Coomassie-stained SDS-Page gels and (B) absorbance measurements showing results of size-exclusion chromatography experiments with Δelbow kinesin-1 tetramers (KHC(Δelbow)/KLC(wt)). BS3 lanes show the mobility of duplicate samples after cross-linking. Complexes eluted in a single peak after the void corresponding to the position of wildtype peak 1. Data are representative of at least 3 independent experiments. (C) Representative electron micrograph showing negative-stained, cross-linked Δelbow/wt sample. Scale bar is 100 nm. Purple boxed particles are expanded below. (D) Immunofluorescence analysis of transfected HeLa cells expressing low levels of HA-Kif5c. Purple arrows show accumulations of Δelbow and Δtail proteins in the cell periphery. Scale bar is 25 μm. Graph shows quantification of percentage of cells with prominent KHC accumulations in the cell periphery from 3 experiments (minimum 25 cells per condition per experiment). ** = p<0.01 using t-test to compare to wildtype to mutants. (E) Schematic illustrating previous Hinge 2 folding model and the new lambda particle model from the present study. Coiled coils are labelled as in Fig. 1 and are approximately to scale. The left schematic shows challenge of rationalising the Hinge 2 model with predicted lengths of coiled-coil sequences to form the compact conformer. Middle panel shows the lambda particle model derived from the present study. Dashed blue lines indicate the unstructured KHC carboxy-terminal tails that contain the IAK regulatory motif. Right panels show equivalent models for the extended conformers observed in this study.

We suggest that the folded kinesin-1 conformer should be known as the kinesin-1 lambda particle (uppercase of the Greek letter, symbol **Λ**),after the autoinhibited dynein phi **(φ)** particle (*51*). The coiled coil architecture revealed by the **Λ**-particle challenges the long-standing assumption that the complex folds on Hinge 2 **(Fig. 4E)**, and so provides the requisite molecular framework to discover how binding of cargo and regulatory factors control the interconversion between this, and the active state(s).

## Acknowledgments

We thank Dr Girish R. Mali, Prof. David S. Stephens, Dr Binyam Mogessie, and Prof. Viki Allan for comments on the project and manuscript.

## Author contributions

Investigation—J.F. Weijman, S.K.N. Yadav, K.J. Surridge, J.A. Cross, U. Borucu, J. Mantell J.; Formal Analysis— J.F. Weijman, S.K.N. Yadav, K.J. Surridge, D.N. Woolfson, C. Schaffitzel, M.P. Dodding; Writing—Original Draft—M.P. Dodding; Writing— Review and Editing— All authors.

## Data and materials availability

All data are available in the main text or the supplementary materials.

## Supplementary Materials

Materials and Methods

Figs. S1 to S7

