## Supplementary material for "Molecular architecture of the autoinhibited kinesin-1 lambda particle": Methods and Supplementary Data

**This PDF file includes:**

Materials and Methods
Figs. S1 to S7

### Materials and Methods

Plasmids and reagents

Rat Kif5c, either full-length (corresponding to residues 2-955) was amplified by polymerase chain reaction (PCR) from HA-tagged expression construct (*8*), and ligated into a His-3C cleavage site-pMW bacterial expression vector. Rat Kif5c ΔTail (corresponding to residues 2-914) was amplified by PCR and inserted into HA-pCB6. Mouse KLC1 was amplified by PCR and ligated into pET28a using NdeI/XhoI sites. The N-terminal His-thrombin cleavage tag was removed by site-directed mutagenesis using the following primers: forward, 5'-GAAGGAGATATAACCATGTATGACAACATGTCCACC-3'; reverse, 5'-GTCATACATGGTTATATCTCCTTCTTAAAGTTAAACAAAATTAT-3'. ΔElbow (Kif5cΔ674-691) was generated by site-directed mutagenesis with the following primers: forward, 5'-GCCCAACTGCAGGATGCTGAGG-3, reverse, 5'-CTGCAGTTGGGCCCGGAGCTT-3'. All plasmids were validated by DNA sequencing.

Protein expression and purification

Heterotetrametric kinesin-1 was expressed in BL21(DE3) cells using a two-plasmid system. BL21(DE3) cells, transformed with both KHC and KLC expression plasmids, were used to inoculate 1 L LB cultures supplemented with both ampicillin and kanamycin. Cells were grown with shaking at 37 $^{o}$C until OD reached 0.6-0.8 before the cultures were cooled to 18 $^{o}$C and protein expression induced with 0.3 µM IPTG. Following shaking incubation at 18 °C overnight, cells were harvested by centrifugation at 6,000 × *g*, 4 $^{o}$C, 15 mins and resuspended (10 mL per 1 L of original culture) in 20 mM HEPES pH 7.4, 300 mM NaCl, 40 mM imidazole before being stored at -20 $^{o}$C. Frozen pellets were thawed and resuspended in a buffer consisting of 40 mM Hepes pH 7.4, 500 mM NaCl, 40 mM imidazole, 5 % (v/v) glycerol, 5 mM betamercaptoethanol (βME) with a Roche complete protease inhibitor tablet added. Cells were lysed by sonication, 0.5 s on, 10 s off, 70% amplitude for 7 min 30 s in an ice bath. Lysate was clarified by centrifugation at 35,000 × *g* JA-20 rotor, 40 mins, 4 $^{o}$C. Clarified lysate was filtered (0.45 µm) before being loaded onto a His-Trap (Sigma) column. Column was washed in base buffer and eluted with a gradient of base buffer up to 500 mM imidazole. Eluted protein was concentrated by ultrafiltration in a 10,000 molecular weight cut-off filter (Cyvita), before snap freezing in liquid nitrogen. Proteins were thawed and further purified by size-exclusion chromatography using a Superose6 10/300 column (Cyvita), in 20 mM HEPES pH 7.4, 150 mM NaCl, 1 mM $MgCl_2$, 0.1 mM ADP, 0.5 mM tris(2-carboxyethyl)phosphine (TCEP, Sigma). For negative stain, freshly eluted proteins from size-exclusion were crosslinked with 0.6 mM bissulfosuccinimidyl suberate (BS3) (ThermoFisher) for 30 minutes at room temperature.

Negative stain electron microscopy

Freshly crosslinked proteins were diluted to 0.003 mg/ml in size-exclusion buffer and 5 µl was pipetted onto a freshly glow-discharged grid (300 mesh copper with formvar/carbon support, TAAB) and incubated at RT for 1 minute. Grids were manually blotted and stained in 5 µl of 3% uranyl acetate (UA) for 1 minute. The UA was blotted, washed in 3% UA and a final 3% UA stain applied for 30 s before blotting and allowing to air dry. Micrographs of grids were acquired on a FEI 120kV BioTwin equipped with an FEI Ceta 4k x 4k CCD camera at 49,000 × magnification corresponding to a pixel size of 2.04 Å/pixel. Particle length was measured using Image J from several micrographs for each complex.

Cryo-negative stain EM Data set acquisition

Negative stain grids were prepared as described above followed by cooling the grid liquid nitrogen immediately prior to loading into the microscope. A data set of 3,407 movies was collected on a FEI Talos Arctica transmission electron microscope operated at 200 kV, equipped with a Gatan K2 Summit direct detector and Gatan Quantum GIF energy filter, operated in zero-loss mode with a slit width of 20 eV using the EPU software. Images were acquired at a nominal magnification of 130,000 ×, giving a pixel size of 1.05 Å/pixel. A 100 µm objective aperture was used. Movies were collected in 55 fractions of 200 ms each. The total dose was 61.05 $e^{-}/Å^{2}$. The movies were acquired with defocus values between -1.0 and -2.5 µm.

Image Processing

Image processing was performed in RELION 3.1.2 (*51*). The movies were motion corrected using MotionCorr2.1 (*52*). CTF-estimation was performed using CTFFIND4 (*53*). Motion corrected and CTF-estimated micrographs were visually inspected manually giving 3,378 final micrographs, a representative example of which is shown in Figure S4. 2,008 particles were manually picked from a randomised 5% subset of micrographs. The manually picked particles were subjected to 2D classification, and the outputs were used to auto-pick the same 5% of micrographs yielding 13,633 particles. The particles were subjected to several rounds of 2D and 3D classification. A resulting 3D model was used to autopick the entire dataset. This yielded 303,353 particles which were extracted and binned 5x to 5.32 Å/pixel. These were subjected to further 2D and 3D classification giving a final cleaned dataset of 22,440 particles (depicted by the 2D class averages present in Figure 2). All 22,440 particles were re-extracted, without binning, and used to produce a single 3D model which was subjected to auto-refinement with a final reported resolution of 30.8 Å.

Back projections of AlphaFold2 model were created by first projecting a volume using the MolMap command in UCSF Chimera. The volume was low-pass filtered to 30 Å, and back-projections created with RELION's *'relion_project'* tool.

Cell culture and immunofluorescence

HeLa cells were grown and maintained in high-glucose Dulbecco's modified Eagle's medium (DMEM; Sigma-Aldrich) supplemented with 10% fetal bovine serum (FBS; Sigma-Aldrich) and 1% penicillin/streptomycin (Gibco) at 37°C with 5% $CO_2$. Transfections and immunofluorescence imaging were as previously described (*54*). Mouse anti-HA (HA-7) was from Sigma-Aldrich and Alexa 568–conjugated anti-mouse was from Thermo Fisher Scientific

AlphaFold2 predictions

Predictions were made using the Google Colab Alphafold Notebook (https://colab.research.google.com/github/deepmind/alphafold/blob/main/notebooks/AlphaFold.ipynb) with a Colab Pro+ subscription using default parameters. The algorithm was asked to model a 2:2 heterotetramer comprised of KHC (Kif5C, NP_001101200.1) residues S410-H917 and KLC (KLC1, NP_032476.2) residues T20-S162, or a homodimer of KHC. These boundaries were established by combining insight from Marcoil predictions shown in Figure 1 and alpha helical domain predictions of the monomeric proteins in the Alphafold-EBI structural database. Default parameters were used and the amber relaxation step was enabled. The single output model was downloaded in pdb format and prepared for presentation using Pymol or UCSF Chimera (*55*). Model statistics are as presented by the software with annotations to highlight the relevant components. For historical reasons reflecting available molecular clones that were subsequently used for the experimental work, models are presented using rat KHC and mouse KLC protein sequences. Mouse, rat and human sequences for KHC and KLC are near identical (>98% sequence identity) and equivalent models were obtained using these orthologues.

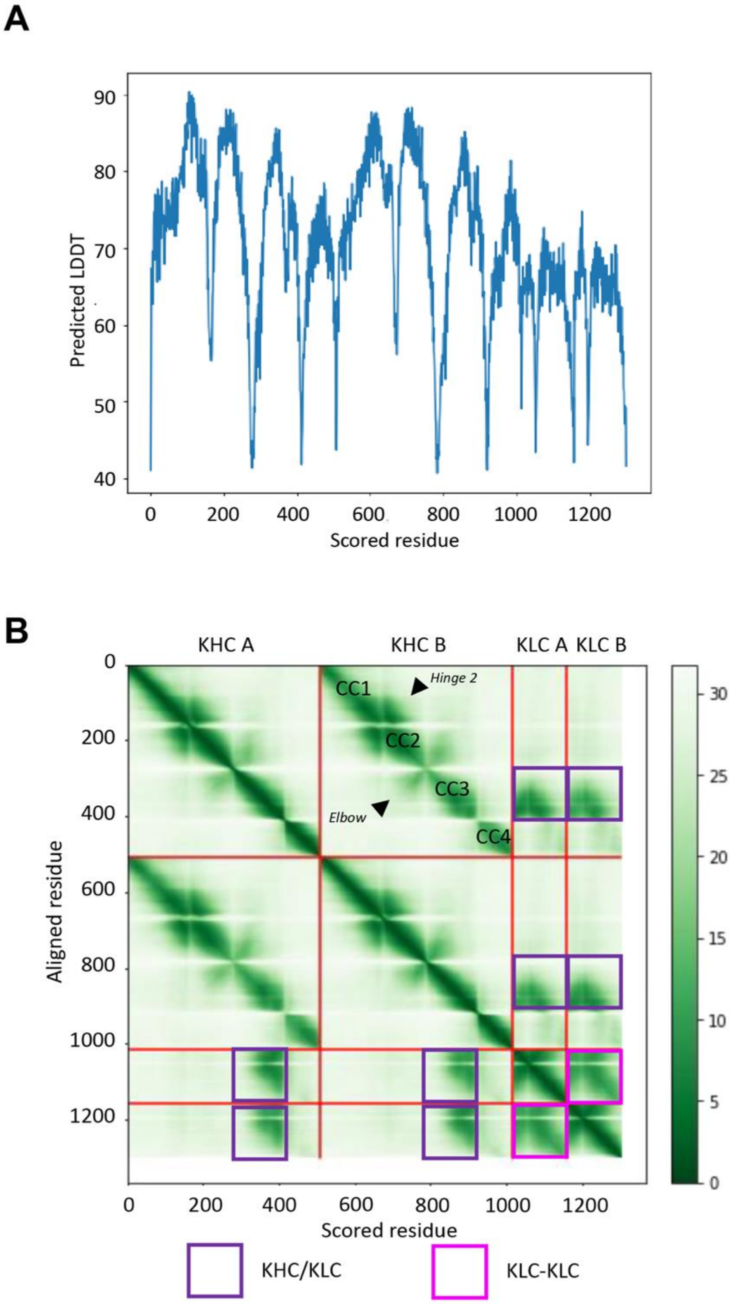


**Fig. S1. AlphaFold2 model statistics for the KHC-KLC heterotetramer** (A) pLDDT (predicted local distance difference test) and (B) predicted aligned error plots for the AlphaFold2-Multimer KHC-KLC model.

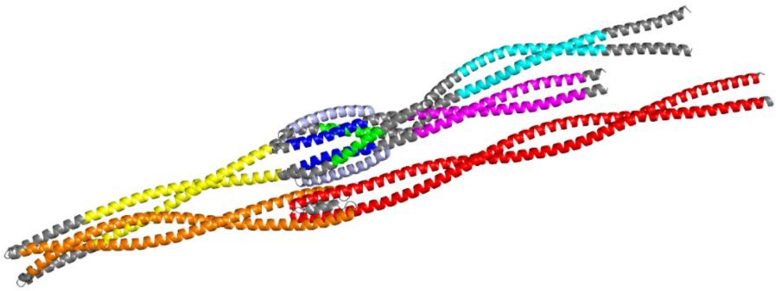

```
KHC Kif5C 410-
SAEKEKYDEEITSLYRQLDDKDDEINQQSQLAEKLKQQMLDQDELLASTRRDYEKIQEELTRLQIENEAAKDEVK
   defgabcdefgabcdefgabcdefgabcdefgabcdefgabcdefgabcdefgabcdefgabcdefgabcde

EVLQALEELAVNYDQKSQEVEDKTRANEQLTDELAQKTTTLTTTQRELSQLQELSNHQKKRATEILNLLLKDLGE
fgabcdefgabcdefgabcdefgabcdefgabcdefgabcdefgabcdefgabcdefgabcdefgabcdefgabc
                                                                 < Tetramer
<------  Hinge 2  ------>
IGGIIGTNDVKTLADVNGVIEEEFTMARLYISKMKSEVKSLVNRSKQLESAQTDSNRKMNASERELAACQLLISQ
defga              defgabcdefgabcdefgabcdefgabcdefgabcdefgabcdefgabcdefgabc
 >                    < Tetramer >
                                                <----- elbow ------>
HEAKIKSLTDYMQNMEQKRRQLEESQDSLSEELAKLRAQEKMHEVSFQDKEKEHLTRLQDAEEVKKALEQQMESH
defgabcdefgabcdefgabcdefgabcdefgabcdefgabcdefga

REAHQKQLSRLRDEIEEKQRIIDEIRDLNQKLQLEQERLSSDYNKLKIEDQEREVKLEKLLLLNDKREQAREDLK
   defgabcdefgabcdefgabcdefgabcdefgabcdefgabcdefgabcdefgabcdefga


GLEETVSRELQTLHNLRKLFVQDLTTRVKKSVELDSDDGGGSAAQKQKISFLENNLEQLTKVHKQLVRDNADLRC
     abcdefgabcdefgabcd                                abcdefgabcdefgabcdef
          AP Trimer

ELPKLEKRLRATAERVKALESALKEAKENAMRDRKRYQQEVDRIKEAVRAKNMARRAH-917
gabcdefgabcdefgabcdefgabcdefga

KLC1 20-
TQDEIISKTKQVIQGLEALKNEHNSILQSLLETLKCLKKDDESNLVEEKSNMIRKSLEMLELGLSEAQ
 abcdefgabcdefgabcdefgabcdefgabcde           abcdefgabcdefgabcdefgab
            AP Trimer                                   AP Trimer

VMMALSNHLNAVESEKQKLRAQVRRLCQENQWLRDELANTQQKLQKSEQSVAQLEEEKKHLEFMNQLK
cd             abcdefgabcdefgabcdefgabcdefgabcdefgabcdefgabcdefgabcd

KYDDDIS-162
```

**Fig. S2. SOCKET2 structure-based assignment of the KHC and KLC heptad register.** Structure shows the AlphaFold2 coiled-coil tetramer prediction in the same orientation as Figure 1C, here coloured by coiled-coil domain predictions from SOCKET2. KHC and KLC sequences are shown below in the same colour scheme with heptad register below. All are predicted parallel coiled coils with exception of the antiparallel (AP) and tetrameric structures highlighted. Sequences are KHC (Kif5C, NP_001101200.1) residues S410-H917 and KLC (KLC1, NP_032476.2) residues T20-S162.

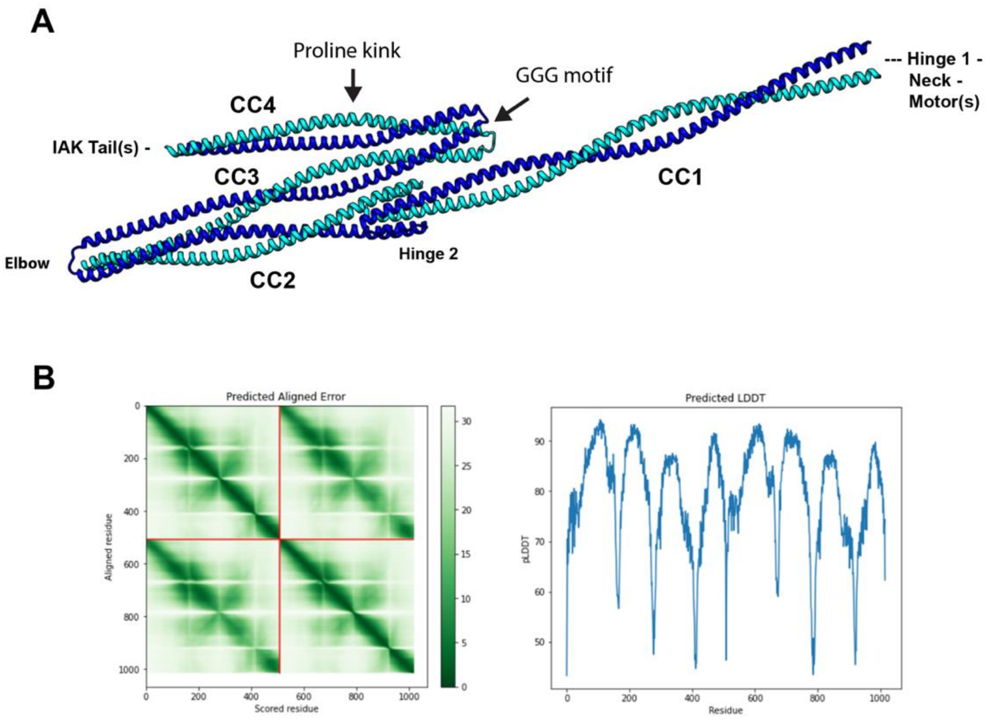


C

```
                                  CC3 (KLC binding)

                                                                        Glycine motif

HsKif5A NP_004975.2         FLYERHEQSKQDLKGLEETVARELQTLHNLRKLFVQDVTTRVKKSAEMEPEDSGGIHSQK  827
HsKif5B NP_004512.1         VMQDRREQARQDLKGLEETVAKELQTLHNLRKLFVQDLATRVKKSAEIDSDDTGGSAAQK  829
HsKif5C NP_004513.1         LLNDKREQAREDLKGLEETVSRELQTLHNLRKLFVQDLTTRVKKSVELDNDDGGGSAAQK  831
RnKif5C NP_001101200.1      LLNDKREQAREDLKGLEETVSRELQTLHNLRKLFVQDLTTRVKKSVELDSDDGGGSAAQK  830
                            .: :::**:::*********::***************::******.*:: :* **  :**

                                  CC4 (Cargo binding)

HsKif5A NP_004975.2         QKISFLENNLEQLTKVHKQLVRDNADLRCELPKLEKRLRATAERVKALEGALKEAKEGAM  887
HsKif5B NP_004512.1         QKISFLENNLEQLTKVHKQLVRDNADLRCELPKLEKRLRATAERVKALESALKEAKENAS  889
HsKif5C NP_004513.1         QKISFLENNLEQLTKVHKQLVRDNADLRCELPKLEKRLRATAERVKALESALKEAKENAM  891
RnKif5C NP_001101200.1      QKISFLENNLEQLTKVHKQLVRDNADLRCELPKLEKRLRATAERVKALESALKEAKENAM  890
                            *************************************************.*******.*

                                               Proline kink
```

**Fig. S3. Structural analysis of KHC** (A) AlphaFold2-Multimer prediction of a homodimer of KHC coiled coils (Kif5C, NP_001101200.1) residues S410-H917. (B) Predicted Aligned Error and pLDDT plots. (C) Clustal Omega multiple sequence alignment from KHC paralogues showing sequences of CC3 and CC4, the di/tri-glycine motif in the linker that separates them, and the proline residue in CC4. Hs – *Homo sapiens*. Rn – *Rattus norvegicus.*

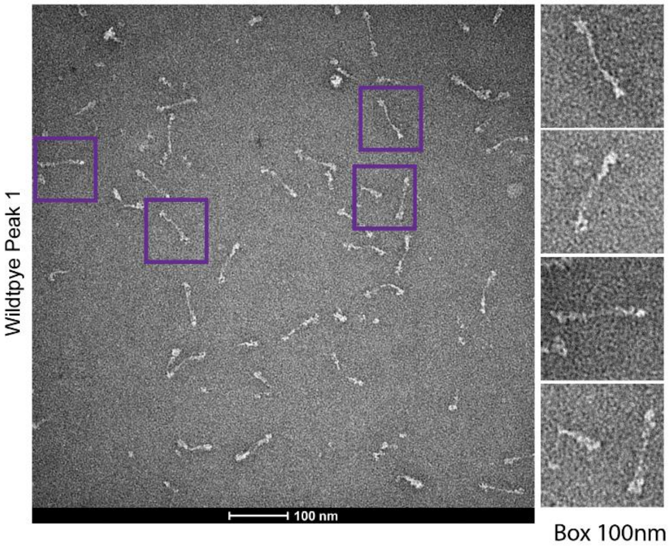


**Fig. S4. Negative stain EM analysis of wildtype peak 1 proteins.** Purple boxes are 100 nm and expanded images of single particles are presented on the right.

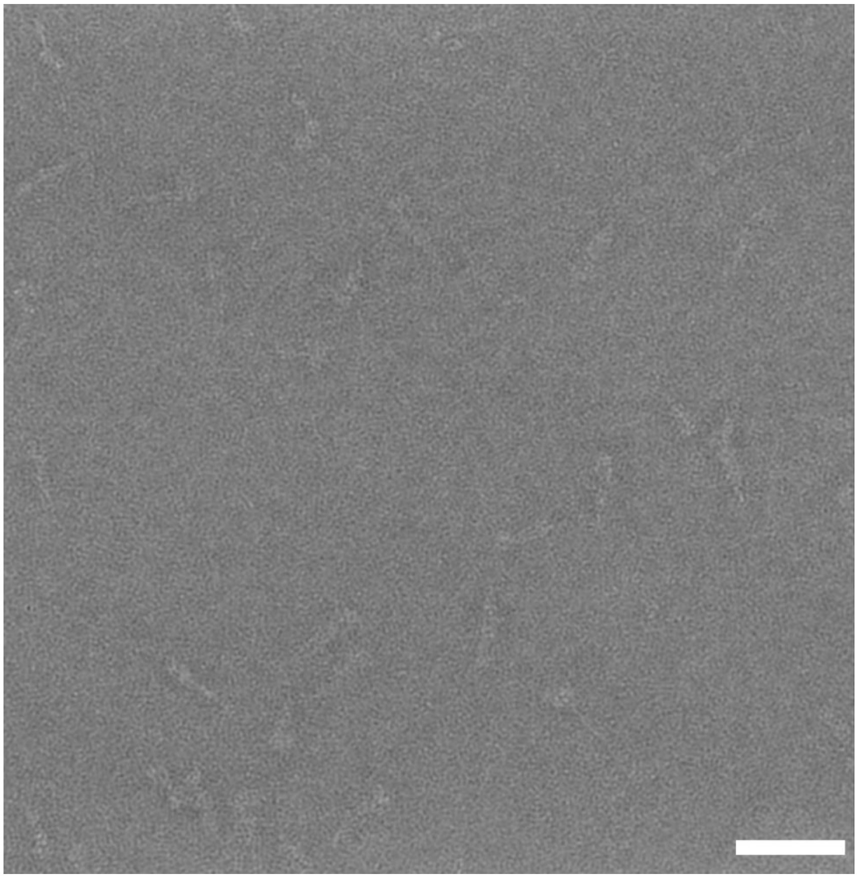

**Fig. S5. Representative cryo-negative stain micrograph.** Example of micrograph used for single particle classification. Scale bar is 50 nm.

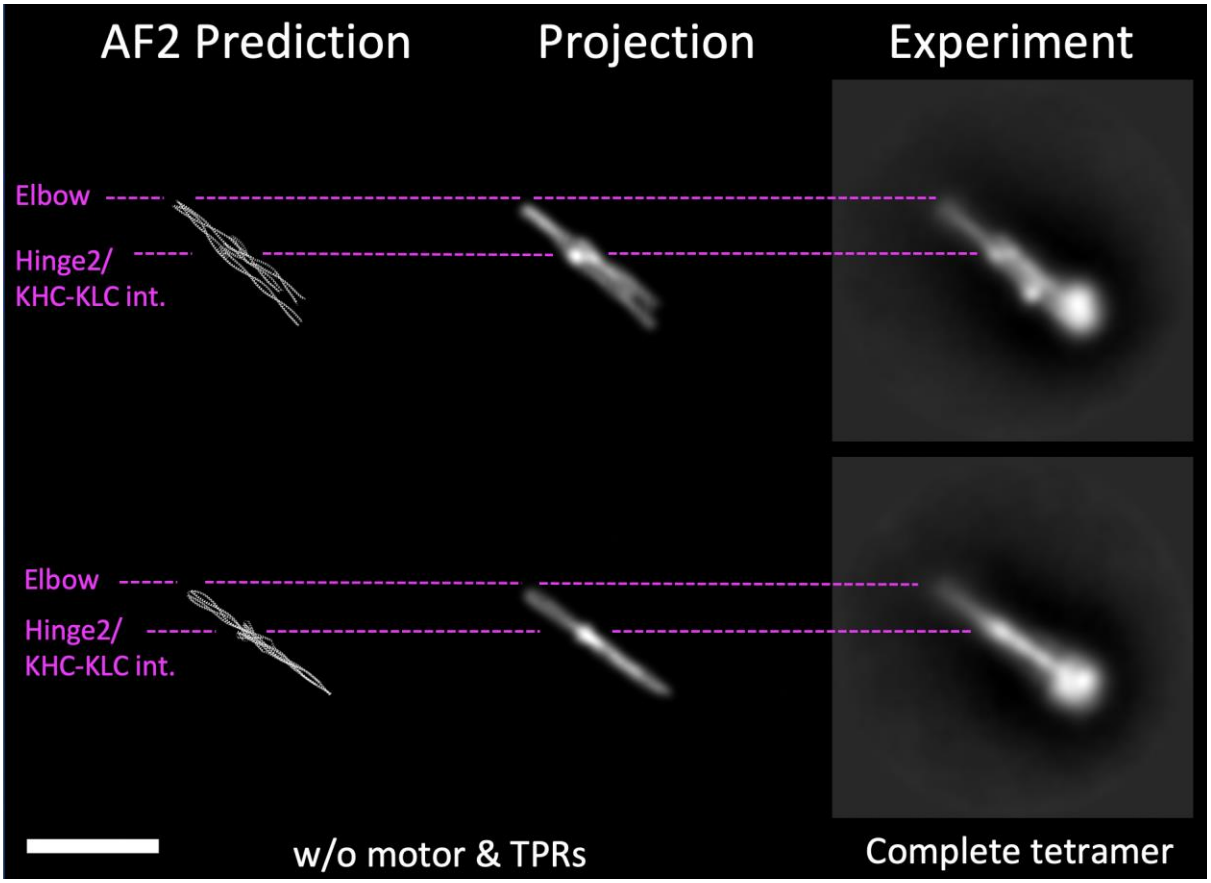


**Fig. S6. Further comparison of computational and experimental data.** Comparison of AlphaFold2 (AF2) atomic model, projection (low pass filtered to 30 Å) with 2D class from experimental data. Scale bar is 25 nm.

```
                         -------Δelbow----- -
651 -       QKRRQLEESQ DSLSEELAKL RAQEKMHEVS FQDKEKEHLT RLQDAEEVKK ALEQQMESHR EAHQKQLSRL RDEIEEKQRI IDEIRDLNQK - 740
MARCOIL     fgabcdefga bcdefgabcd efgabcdebc defgabcdef gabcdefgab cdefgabcde fgaefgabcd efgabcdefg abcdefgabc
SOCKET      fgabcdefga bcdefgabcd efgabcdefg a                                defgabcd efgabcdefg abcdefgabc
AF2         HHHHHHHHHH HHHHHHHHHH HHHHHHHHHH HHHH-----H HHHHHHHHHH HHHHHHHHHH HHHHHHHHHH HHHHHHHHHH HHHHHHHHHH
PSIPRED4    HHHHHHHHHH HHHHHHHHHH HHHHHHHHHH HHHHHHHHHH HH-HHHHHHH HHHHHHHHHH HHHHHHHHHH HHHHHHHHHH HHHHHHHHHH
JPRED4      HHHHHHHHHH HHHHHHHHHH HHHHHHHHHH HHHHHHHHHH ----HHHHHH HHHHHHHH-H HHHHHHHHHH HHHHHHHHHH HHHHHHHHHH

            LKKRHLEESY DSLSDELAKL QAQETVHEVA LKDKEP---- DTQDADEVKK ALELQMESHR 707        Kif5A Hs NP_004975.2
            QKKRQLEESV DALSEELVQL RAQEKVHEM- ----EKEHLN KVQTANEVKQ AVEQQIQSHR 709        Kif5B Hs NP_004512.1
            QKRRQLEESQ DSLSEELAKL RAQEKMHEVS FQDKEKEHLT RLQDAEEMKK ALEQQMESHR 711        Kif5C Hs NP_004513.1
            QKRRQLEESQ DSLSEELAKL RAQEKMHEVS FQDKEKEHLT RLQDAEEVKK ALEQQMESHR 710        Kif5C Rn NP_001101200.1
             *:*:****  *:**:**.:* :***.:**:      *         * *:*:*: *:* *::***
```

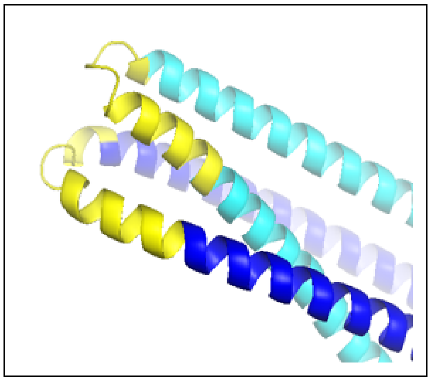

**Fig. S7. Bioinformatic analysis of the kinesin-1 elbow.** Alignment showing primary sequence of the Kif5C elbow region with Marcoil heptad prediction below in rainbow colours. The elbow deletion is highlighted in yellow on the sequence and also on the AlphaFold2 structure presented below. Secondary structure predictions from the AlphaFold2 model (AF2), Psipred4 and Jpred4 are also shown, along with a Clustal Omega multiple sequence alignment highlighting conservation and divergence of residues across this region in the KHC paralogues. Hs – *Homo sapiens*. Rn – *Rattus norvegicus.*
